# FINEMAP: Efficient variable selection using summary data from genome-wide association studies

**DOI:** 10.1101/027342

**Authors:** Christian Benner, Chris C.A. Spencer, Samuli Ripatti, Matti Pirinen

## Abstract

**Motivation:** The goal of fine-mapping in genomic regions associated with complex diseases and traits is to identify causal variants that point to molecular mechanisms behind the associations. Recent fine-mapping methods using summary data from genome-wide association studies rely on exhaustive search through all possible causal configurations, which is computationally expensive.

**Results:** We introduce FINEMAP, a software package to efficiently explore a set of the most important causal configurations of the region via a shotgun stochastic search algorithm. We show that FINEMAP produces accurate results in a fraction of processing time of existing approaches and is therefore a promising tool for analyzing growing amounts of data produced in genome-wide association studies.

**Availability:** FINEMAP v1.0 is freely available for Mac OS X and Linux at http://www.christianbenner.com.

## 1 Introduction

Genome-Wide Association Studies (GWAS) have identified thousands of genomic regions associated with complex diseases and traits. Any associated region may contain thousands of genetic variants with complex correlation structure. Therefore, one of the next challenges is *fine-mapping* that aims to pinpoint individual variants and genes that have a direct effect on the trait. This step is crucial for fully exploiting the potential of GWAS: to unveil molecular biology of complex traits and, eventually, provide targets for therapeutic interventions. For a recent review on fine-mapping, see Spain *et al.* (2015).

A standard approach for refining association signals is a step-wise conditional analysis, an iterative procedure that conditions on the Single-Nucleotide Polymorphisms (SNPs) with the lowest P-value of association until no additional SNP reaches the pre-assigned P-value threshold. While conditional analysis is informative about the number of complementary sources of association signals within the region, it fails to provide probabilistic measures of causality for individual variants. To overcome this problem, many recent fine-mapping methods have adopted a Bayesian framework.

Approaches for Bayesian analysis of multi-SNP GWAS data include exhaustive search as implemented in software BIMBAM (Servin *et al.*, 2007), MCMC algorithms (Guan *et al.*, 2011), variational approximations (Carbonetto *et al.*, 2012) and stochastic search as implemented in software GUESS (Bottolo *et al.*, 2010, 2013) and GUESSFM (Wallace *et al.*, 2015). Bayesian fine-mapping has also been conducted under a simplified assumption of a single causal variant in the region (WTCCC *et al.*, 2012). Common to these approaches is that they require original genotype-phenotype data as input, which is becoming impractical or even impossible as the size of current GWAS meta-analyses rises to several hundreds of thousands of samples (Wood *et al.*, 2014). For this reason, fine-mapping methods have recently been extended to use only GWAS summary data together with a SNP correlation estimate from a reference panel. To our knowledge, the existing fine-mapping implementations using GWAS summary data are PAINTOR (Kichaev *et al.*, 2014, 2015), CAVIAR (Hormozdiari *et al.*, 2014) and CAVIARBF (Chen *et al.*, 2015).

PAINTOR is an EM-algorithm to jointly fine-map several associated regions by utilizing functional annotation information of individual variants. As a special case of only a single region without annotation information, PAINTOR tackles the standard fine-mapping problem. CAVIAR differs from PAINTOR by modeling the uncertainty in the observed association statistics. This might be a reason why CAVIARBF, a more efficient implementation of CAVIAR, has been reported to be more accurate than PAINTOR in prioritizing variants when no annotation information is available (Chen *et al.*, 2015).

Although PAINTOR, CAVIAR and CAVIARBF are very useful methods for performing fine-mapping on GWAS summary data, we think that their implementation via an exhaustive search through all possible causal configurations is likely to hinder their use in several settings. For example, it becomes computationally slow or even impossible to run these methods by allowing more than three causal variants on dense genotype data with thousands of variants per region. Thus, these methods are unlikely to make full use of unprecedented statistical power to discern complex association patterns provided by ever increasing GWAS sample sizes and genome sequencing technologies.

We introduce FINEMAP, a novel software package to improve the performance of GWAS summary data based fine-mapping. The statistical model of FINEMAP is similar to CAVIAR and CAVIARBF while the important difference is the computational algorithm. FINEMAP uses a Shotgun Stochastic Search (SSS) algorithm (Hans *et al.*, 2007) that explores the vast space of causal configurations by concentrating efforts on the configurations with non-negligible probability. We compare FINEMAP with the exhaustive search algorithm implemented in CAVIARBF. The comparisons to two other GWAS summary statistics based fine-mapping methods CAVIAR and PAINTOR are not shown in this paper since CAVIARBF is more efficient but equally accurate as CAVIAR and more accurate than PAINTOR without annotation information (Chen *et al.*, 2015). In this paper we show that

- FINEMAP is thousands of times faster than CAVIARBF while still providing similar accuracy in the examples where CAVIARBF can be applied.
- FINEMAP is more accurate than CAVIARBF when the number of causal variants in CAVIARBF needs to be restricted for computational reasons.

Our examples are based on genotype data of the Finnish population as well as summary statistics from GWAS on Parkinson’s disease (UKPDC and WTCCC2, 2011).

## 2 Methods

We are interested in fine-mapping a genomic region using GWAS summary data instead of original genotype-phenotype data as input. The building blocks of our Bayesian approach are the likelihood function (subsection 2.1), priors (subsection 2.2), efficient likelihood evaluation (subsection 2.3) and efficient search algorithm (section 3). At each step we describe how our choices differ from the existing methods PAINTOR and CAVIARBF.

### 2.1 Likelihood function

For a quantitative trait, we assume the following linear model

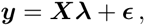

where ***y*** is a mean-centered vector of values of a quantitative trait for *n* individuals, ***X*** a column-standardized SNP genotype matrix of dimension *n* × *m* and 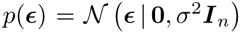. The Maximum Likelihood Estimate (MLE) of the causal SNP effects **λ** depends on ***X*** and ***y*** only through the SNP correlation matrix ***R*** = *n*^−1^***X****^T^****X*** and single-SNP *z*-scores *ẑ* = *(nσ*^2^)^−1/2^***X***^T^***y***

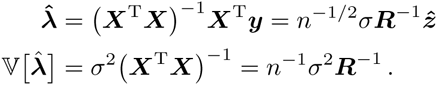

Thus, it is possible to approximate the likelihood function for **λ** by 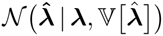 using a SNP correlation estimate from a reference panel and single-SNP *z*-scores from a standard GWAS software, as previously done in GCTA (Yang *et al.*, 2011), PAINTOR and CAVIARBF. For binary traits, a similar approximation applies with *z*-scores originating from logistic regression and *σ*^2^ ≈ 1/{*ϕ*(1 − *ϕ*)}, where *ϕ* is the proportion of cases among the *n* individuals (Pirinen *et al.*, 2013).

When *m* is large but **λ** has only very few non-zero elements, the MLE alone is not ideal since it does not account for the sparsity assumption (Figure 1). Thus, we take a Bayesian approach with a prior distribution that induces sparsity among causal effects.

**Figure 1:**
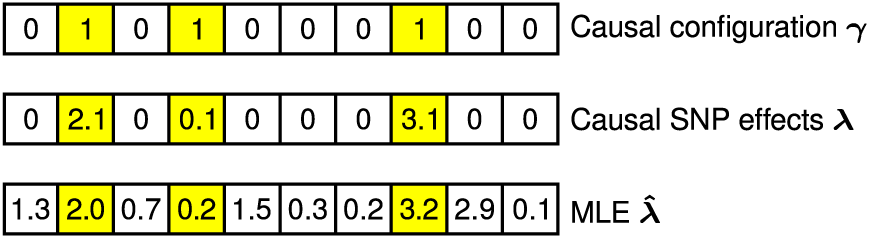
The binary indicator vector *γ* determines which SNPs have non-zero causal effects 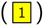. The underlying causal (linear) model for a quantitative trait assumes only few SNPs with a causal effect. The Maximum Likelihood Estimate (MLE) of the causal SNP effects 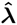 can be computed by using only the SNP correlation matrix and single-SNP *z*-scores. However, the MLE is not ideal because it does not account for the sparsity assumption.

### 2.2 Priors for λ and γ

Let a binary indicator vector **γ** determine which SNPs have non-zero causal effects (*γ_ℓ_* = 1 if the *ℓ*th SNP is causal and 0, otherwise; see top panel in Figure 1). For the causal effects, we use the prior

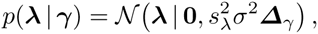

where 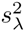 is the prior variance for the causal effects in units of *σ*^2^ and **Δ***_γ_* a diagonal matrix with *γ* on the diagonal. In our examples for quantitative traits, we have set 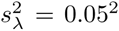 and *σ*^2^ to the observed variance of the trait. This means that with 95% probability a causal SNP explains less than 1% of the trait variation. When available *z*-scores originate from logistic regression, we have set the product 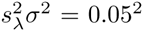. This means that with 95% probability the effect of a causal SNP on the odds-ratio scale is less than 1.15 for common variants (MAF = 0.5) and less than 2.0 for low-frequency variants (MAF = 0.01), where MAF is the minor allele frequency.

To define the prior for each causal configuration, we use a general discrete distribution for the number of causal SNPs

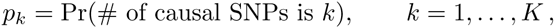

where *K ≪ m* is the maximum number of SNPs in the causal configuration. Note that we assume that the region to be fine-mapped includes at least one causal SNP, i.e., *p*_0_ = 0. For a fixed value of *k*, we assume the same probability for each configuration with *k* causal SNPs. Thus, a priori,

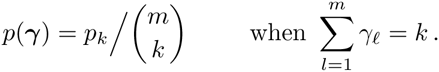

PAINTOR does not use an explicit prior on *k* but restricts *k* ≤ 3 in practice. The default prior used by CAVIARBF assumes that each SNP is causal with probability 1/*m* and that *k* ≤ 5. This is a special case of our prior when we set

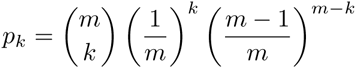

and renormalize for *K* = 5 except that CAVIARBF assigns non-zero prior also for the null configuration *k* = 0.

### 2.3 Marginal likelihood for γ

We now show how the marginal likelihood for the causal configuration **γ** can be computed efficiently.

#### 2.3.1 Integrating out causal effects A

The likelihood function *p*(**y** *|* **λ**, ***X***) of the causal SNP effects is (proportional to) a Normal density 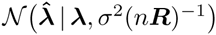 This enables an analytic solution for the marginal likelihood of **γ** eliminating the causal effects

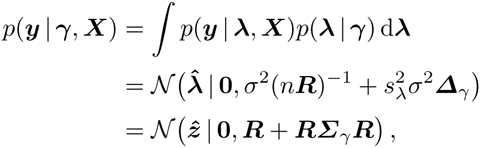

where we defined 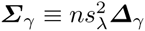. Importantly, an evaluation of the marginal likelihood requires only single-SNP *z*-scores and SNP correlations from a reference panel. This elimination of **λ** is similar to the one used by CAVIARBF and differs from PAINTOR that fixes those values based on the observed *z*-scores. Next, we describe two implementations to evaluate 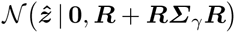 with high computational efficiency.

#### 2.3.2 Reducing the complexity from 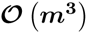 to 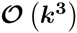

##### Option 1

Let *C* = {1,…, *k}* and *N* = *{k* + 1,…, *m}* be respectively the set of causal and non-causal SNPs. Consider the quadratic form

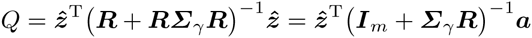

inside the exponential function in 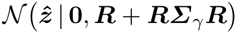, where *a* = *R*^−1^ *ẑ* can be precomputed. We solve the linear system (***I****_m_* + ***Σ_γ_R***)***b*** = ***a*** for ***b*** by observing that the *m − k* elements in ***b*** corresponding to non-causal SNPs (*γ_ℓ_* = 0) are *b_ℓ_* = *a_ℓ_* and the remaining elements result from solving a system of *k* equations

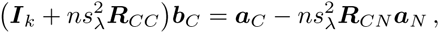

where ***R****_CC_* is the *k* × *k* correlation matrix of the causal SNPs and *R_CN_* the *k × (m − k)* submatrix of ***R*** corresponding to the cross-covariances between the causal and non-causal SNPs. In addition, we observe that det(***I****_m_* + ***Σ****_γ_R*) is simply det 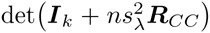 after expanding with respect to the rows corresponding to non-causal SNPs. Computationally, these computations require one Cholesky decomposition with complexity 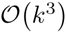 and provide thus a considerable saving compared to the naive way of decomposing the whole *m* × *m* matrix with complexity 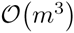.

This derivation differs from the one used by CAVIARBF that is similar to our option 2 below. It also differs from PAINTOR that fixes **λ** based on the observed *z*-scores and performs once a Choleksy decomposition of the whole *m* × *m* SNP correlation matrix that is used repeatedly in each likelihood evaluation.

##### Option 2

We partition the observed *z*-scores into components ***ẑ****_C_* and ***ẑ****_N_* and permute rows and columns of the SNP correlation matrix and covariance matrix *Σ_γ_* such that

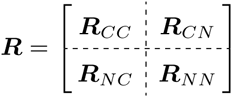

and *Σ_γ_* = diag{*σ_ℓ_*} with *σ_k_*_+1_ = ⋯ = *σ_m_* = 0. This partitioning entails a block structure in the covariance matrix of 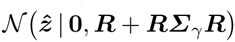

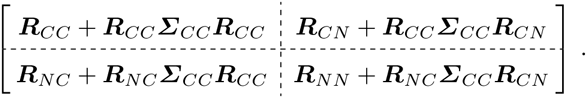

Using properties of the multivariate Normal distribution, the conditional expectation and covariance matrix of *ẑ_N_* given *ẑ_C_* are readily available

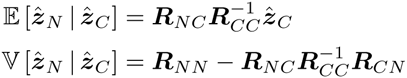

and do not dependent on ***Σ****_γ_*. We rewrite the marginal likelihood 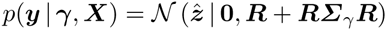 in terms of the marginal distribution of *ẑ_C_* and conditional distribution of *ẑ_N_* given *ẑ_C_* to obtain the following expression

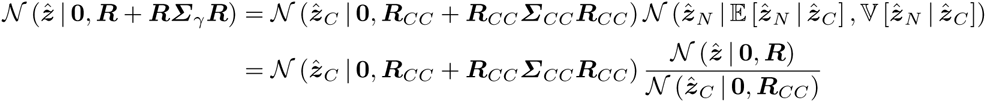

This means that we can compute the Bayes factor for assessing the evidence against the null model by using only the causal SNPs

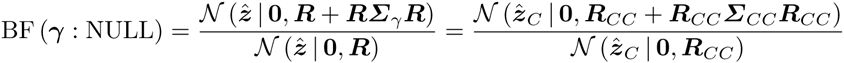

and that the marginal likelihood is proportional to this expression. CAVIARBF utilizes this result, although without a mathematical derivation explicitly shown in Chen *et al.* (2015).

### 2.4 Posterior for γ

According to the Bayesian paradigm, we want to base our inference on the posterior of causal configurations *p(****γ*** | ***y***, ***X***) The unnormalized posterior can be evaluated by combining the prior with the marginal likelihood (option 1) as

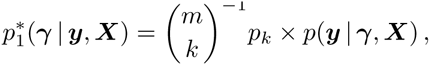

where *k* is the number of causal SNPs in configuration ***γ***. In addition, we can compute unnormalized posterior by using the Bayes factor (option 2)

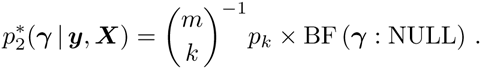

We observed that option 2 was faster than option 1 and therefore option 2 is used by default in FINEMAP. Ideally, *p*(****γ*** *|* ***y***, ***X***) were normalized over all 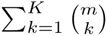 causal configurations. Unfortunately, this is computationally intractable already for modest values of *K* > 5. However, as we show in the results section, typically a large majority of the causal configurations have negligible posterior probability and hence a good approximation for the posterior can be achieved by concentrating on only those with non-negligible probability. We explore the space of causal configurations with a Shotgun Stochastic Search (SSS) algorithm (Hans *et al.*, 2007) that rapidly evaluates many configurations and is designed to discover especially those with highest posterior probability.

## 3 Shotgun stochastic search

We use SSS to efficiently evaluate many causal configurations and discover especially those with highest posterior probability. SSS conducts a pre-defined number of iterations within the space of causal configurations. In each iteration (Figure 2), the neighborhood of the current causal configuration is defined by configurations that result from deleting, changing or adding a causal SNP from the current configuration. The next iteration starts by sampling a new causal configuration from the neighborhood based on *p*(****γ*** | ***y***, ***X***) normalized within the neighborhood. All evaluated causal configurations and their unnormalized posterior probabilities are saved in a list Γ* for downstream analyses. The aim of the algorithm is that Γ* contains all relevant causal configurations, that is, those with non-negligible posterior probabilities.

**Figure 2:**
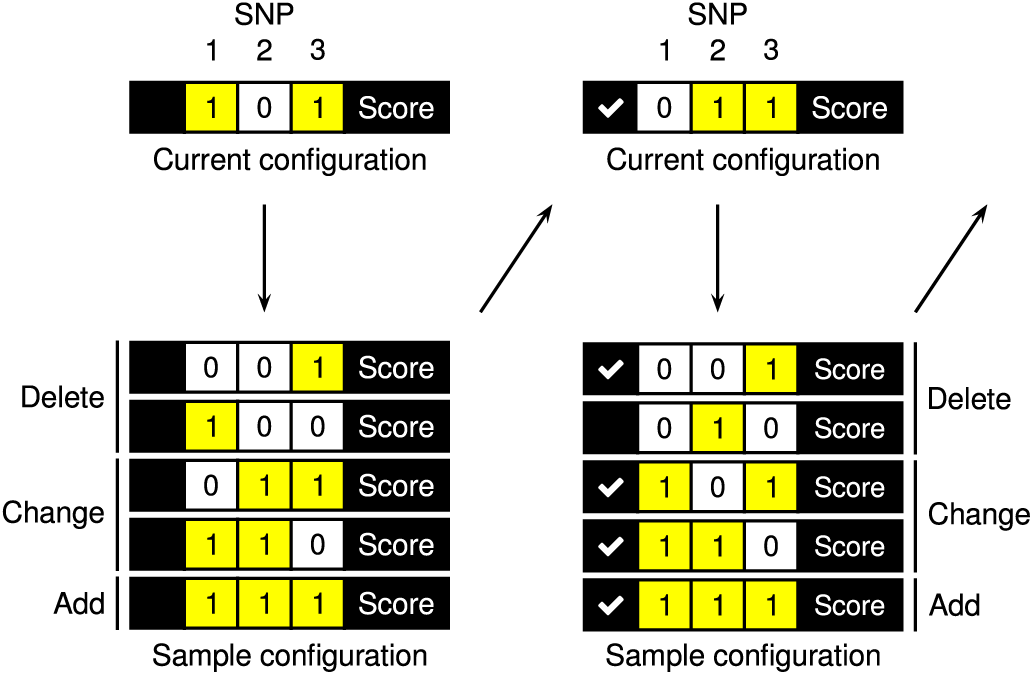
Shotgun stochastic search rapidly identifies configurations of causal SNPs with high posterior probability. In each iteration, the neighborhood of the current causal configuration is defined by configurations that result from deleting, changing or adding a causal SNP 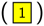 from the current configuration. The next iteration starts by sampling a new causal configuration from the neighborhood based on the scores normalized within the neighborhood. The unnormalized posterior probabilities remain fixed throughout the algorithm and can thus be memorized (✓) to avoid recomputation when already evaluated configurations appear in another neighborhood.

The posterior probability that SNPs in configuration ***γ*** are causal is computed by normalizing over Γ*

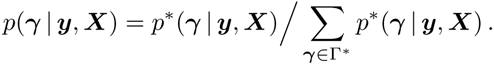

We compute the marginal posterior probability that the *ℓ*th SNP is causal, also called single-SNP inclusion probability, by averaging over all evaluated configurations

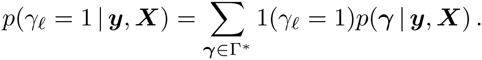

In addition, we compute a single-SNP Bayes factor for assessing the evidence that the *ℓ*th SNP is causal as

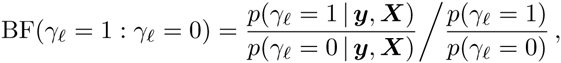

where the prior probability of the *ℓ*th SNP being causal is

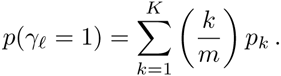

PAINTOR, CAVIAR and CAVIARBF do not perform a stochastic search but enumerate all causal configurations with *k* = 1, … *, K*. When *m* is large but there are only few true causal SNPs, the exhaustive search is computationally expensive and inefficient since most configurations make a negligible contribution to the single-SNP inclusion probabilities.

### 3.1 Computational implementation

For 1 *< k < K*, the number of causal configurations to be evaluated in each iteration is:

- *k* for deleting,
- *k*(*m* − *k*) for changing,
- *m − k* for adding a causal SNP

Computing *p*(****γ*** | ***y***, ***X***) requires a Cholesky decomposition with complexity 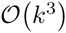 that is fast when *K ≪ m.* Importantly, each unnormalized posterior probability remains fixed throughout the algorithm. This means that we can use a hash table (std::unordered_map in C++) to avoid recomputing *p*(****γ*** | ***y***, ***X***) when already-evaluated configurations appear in another neighborhood. Inserting to and retrieving from the hash table requires constant time on average. Hash table lookups reduce the dominant computational cost of the algorithm: exploring the vast space of causal configurations. This renders SSS computational efficient because it traverses the space of causal configurations by moving back and forth to configurations with high posterior probability and overlapping neighborhoods.

## 4 Test data generation

We obtained real genotype data on 18834 individuals from the National FINRISK study (Vartiainen *et al.*, 2010). The genotype data comprise a 500 kilobase region centered on rs11591147 in *PCSK9* gene on chromosome 1 with 1920 polymorphic SNPs with pairwise absolute correlations less than 0.99. To assess the computational efficiency and fine-mapping accuracy, we considered the following scenarios:

- **Scenario A** Increasing number of SNPs (*m* = 750,1000,1250,1500) considering causal configurations with up to *K* = 3 or *K* = 5 SNPs.
- **Scenario B** Fixed number of *m* = 150 SNPs considering causal configurations with increasing maximum number of SNPs (*K* = 1, 2, 3,4, 5).

We generated data sets where causal SNPs had highly correlated proxies since this is a setting where an in-exhaustive search could theoretically have problems. Five hundred data sets were generated under each combination of *m* and *K* in scenarios A and B using the following linear model:

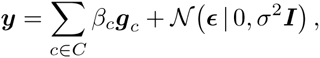

where *C* is the set of causal SNPs, ***g****_c_* the vector of genotypes at the *c*th causal SNP, *β_c_* and *f_c_* respectively the effect size and minor allele frequency of the *c*th causal SNP and 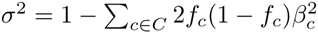. The number of causal SNPs was five in scenario A and *K* = 1, …, 5 in scenario B. In each data set, the causal SNPs were randomly chosen among those variants that had highly correlated proxies (absolute correlation greater than 0.5) among the other variants. The effect sizes of the causal SNPs were specified so that the statistical power at a significance level of 5 × 10^−8^ was approximately 0.5. Single-SNP testing using a linear model was performed to compute *z*-scores. Each set of *z*-scores was then analyzed with CAVIARBF (default parameters) and FINEMAP (100 iterations saving the top 50000 evaluated causal configurations). For both methods, the prior standard deviation of the causal effects was set to 0.05 and the prior distribution of each configuration with *k* causal SNPs was specified as

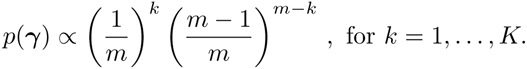

This required excluding the null configuration (*k* = 0) from the output of CAVIARBF.

## 5 Results

The main difference between FINEMAP and CAVIARBF is the search strategy to explore the space of causal configurations. We compare the computational efficiency and fine-mapping accuracy of FINEMAP with CAVIARBF to assess the impact of replacing exhaustive with stochastic search. We also illustrate FINEMAP on data from the UK Parkinson’s Disease Consortium and the Wellcome Trust Case Control Consortium 2 by fine-mapping 4q22/SNCA region that contains a complex association pattern with Parkinson’s disease (The UKPDC and WTCCC2, 2011).

### 5.1 Computational efficiency

The left panel of Figure 3 shows that FINEMAP is thousands of times faster than CAVIARBF when considering causal configurations with up to three SNPs in Scenario A. The difference in processing time becomes even larger when the maximum number of possible causal SNPs increases (Scenario B) in the right panel of Figure 3. CAVIARBF slows down quickly due to the exhaustive search but FINEMAP’s processing time does not increase considerably with increasing *K*. Importantly, there is no need to restrict the number of causal SNPs in FINEMAP to small values *(K* ≤ 5) as is necessary for CAVIARBF.

**Figure 3:**
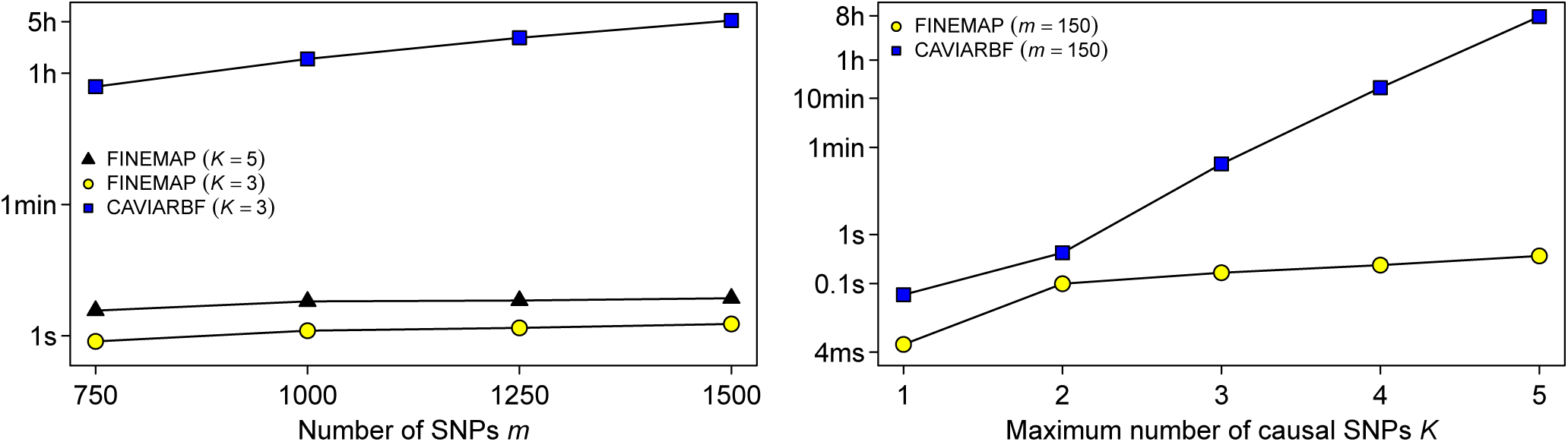
Processing time of one locus with FINEMAP and CAVIARBF on logio scale. Left panel: Scenario A with increasing number of SNPs allowing *K* = 3 or *K* = 5 causal SNPs. Right panel: Scenario B with 150 SNPs considering causal configurations with different maximum numbers of SNPs. All processing times are averaged over 500 data sets using one core of a Intel Haswell E5–2690v3 processor running at 2.6GHz.

### 5.2 Fine-mapping accuracy

We computed the maximum absolute differences between the single-SNP inclusion probabilities in each data set under scenario B to assess the fine-mapping accuracy of FINEMAP and CAVIARBF (Table 1). The small differences (max < 0.11, median < 6 × 10^−4^) show that for practical purposes FINEMAP achieves similar accuracy as CAVIARBF despite concentrating only on a small but relevant subset of all possible causal configurations (see Discussion). Figure 4 shows details of those SNPs in Scenario B for which the difference between the methods is larger than 0.01. We see that by ignoring the large majority of very improbable configurations, FINEMAP slightly overestimates the largest probabilities, that typically belong to the truly causal SNPs, and underestimates smaller probabilities, that most often belong to the non-causal SNPs.

**Figure 4:**
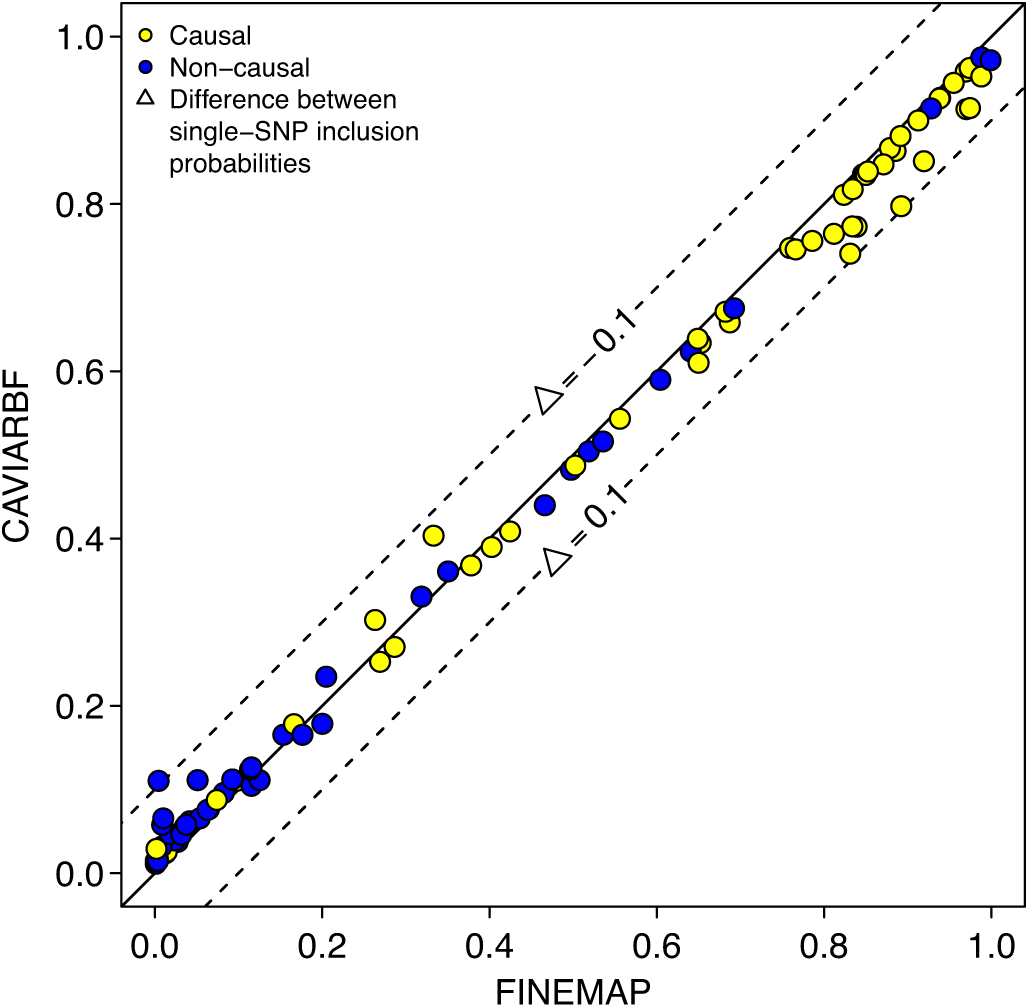
Single-SNP inclusion probabilities of all SNPs in Scenario B with absolute difference larger than 0.01 between FINEMAP and CAVIARBF.

**Table 1.** Percentiles of absolute maximum differences between FINEMAP’s and CAVIARBF’s single-SNP inclusion probabilities in Scenario B

| $m = 150 \mid K$ | 1 | 2 | 3 | 4 | 5<sup>a</sup> |
| --- | --- | --- | --- | --- | --- |
| Max | 5e-7 | 8e-3 | 2e-2 | 1e-1 | - |
| 99th percentile | 4e-7 | 2e-3 | 8e-3 | 4e-2 | - |
| 95th percentile | 3e-7 | 5e-4 | 3e-3 | 1e-2 | - |
| Median | 4e-8 | 4e-7 | 2e-5 | 6e-4 | - |

<sup>a</sup>CAVIARBF could not compute single-SNP inclusion probabilities due to a memory allocation failure (std::bad\_alloc).

In addition to considering only causal configurations with up to three SNPs under scenario A, we also ran FINEMAP with *K* = 5 to demonstrate the increase in fine-mapping performance in this case where the true number of causal SNPs was five. We determined the proportion of causal SNPs that are included when selecting different numbers of top SNPs on the basis of ranked single-SNP inclusion probabilities (Figure 5). FINEMAP and CAVIARBF had the same performance when considering causal configurations with up to three SNPs in genomic regions with 1500 SNPs. (Similar performance was also observed for genomic regions with different numbers of SNPs.) As expected, FINEMAP showed better fine-mapping performance when considering causal configurations with up to five SNPs.

**Figure 5:**
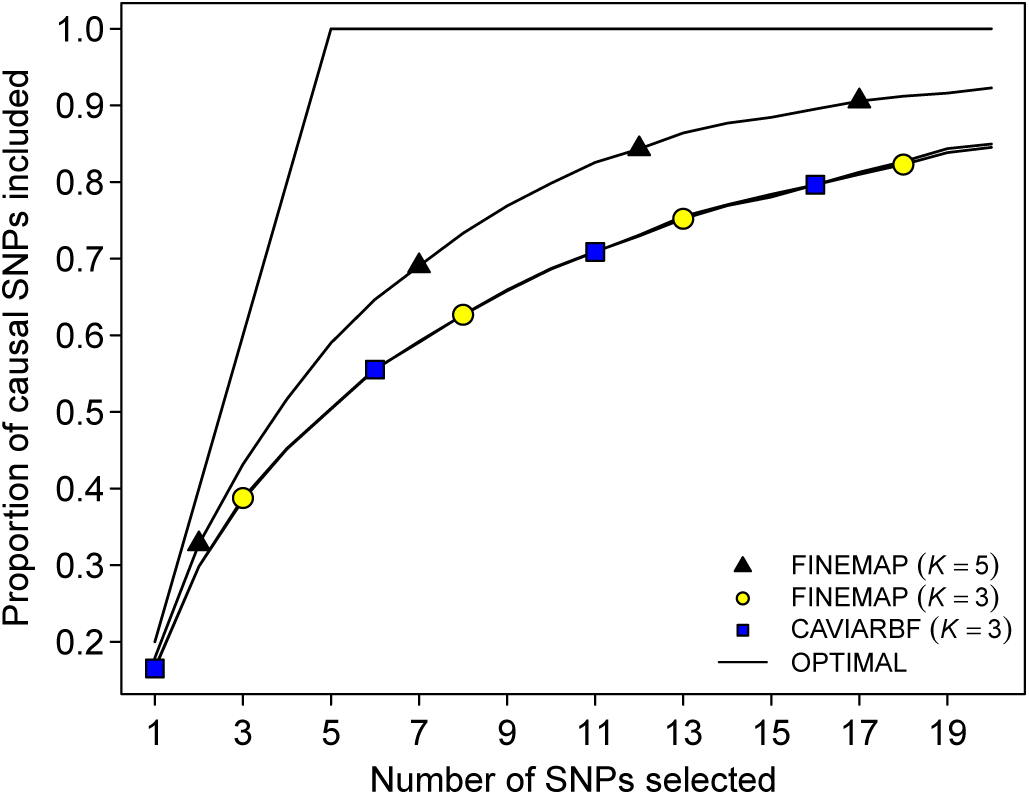
Fine-mapping accuracy of FINEMAP and CAVIARBF on data with five causal SNPs, allowing either *K* = 3 or *K* = 5 causal SNPs. The proportion of causal SNPs included is plotted against the number of top SNPs selected on the basis of ranked single-SNP inclusion probabilities. Proportions are averaged over 500 data sets with 1500 SNPs. Case *K* = 5 is computationally intractable for CAVIARBF.

### 5.3 4q22/SNCA association with Parkinson’s disease

Using single-SNP testing, the UKPDC and WTCCC2 (2011) found evidence for an association with Parkinson’s disease in the 4q22 region with the lowest P-value at rs356220. A conditional analysis on rs356220 revealed a second SNP rs7687945 with P-value 3 × 10^−5^ that in the single-SNP testing had only a modest P-value of 0.13. These two SNPs are in low Linkage Disequilibrium (LD) (*r*^2^ = 0.168 in the original data) but the LD was sufficient enough to mask the effect of rs7687945 in single-SNP testing. This complex pattern of association was replicated in an independent French data set (UKPDC and WTCCC2, 2011).

To test whether FINEMAP is able to pick up this complex association pattern, we extracted a 2 megabase region centered on rs356220 with 363 directly genotyped SNPs from the original genotype data. Single-SNP testing using a logistic model implemented in SNPTEST was performed to compute *z*-scores. The dataset was then analyzed with FINEMAP using 100 iterations and prior parameter value of 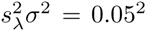. Top panel of Figure 6 shows that the evidence that rs356220 and rs7687945 are causal is the largest among all SNPs. In addition, the causal configuration that simultaneously contains both rs356220 and rs7687945 has the highest posterior probability (0.132). The second most probable (0.113) causal configuration contains rs356220 and rs2301134. High correlation between rs7687945 and rs2301134 (*r*^2^ = 0.974) explains why these two SNPs are difficult to tell apart. We conclude that FINEMAP was able to identify the complex association pattern at the second SNP that only became identifiable after the first SNP was included in the model. As opposed to the standard conditional analysis, FINEMAP provides posterior probabilities for all SNPs in the region and is thus able to simultaneously identify many causal variants without a step-wise procedure.

**Figure 6:**
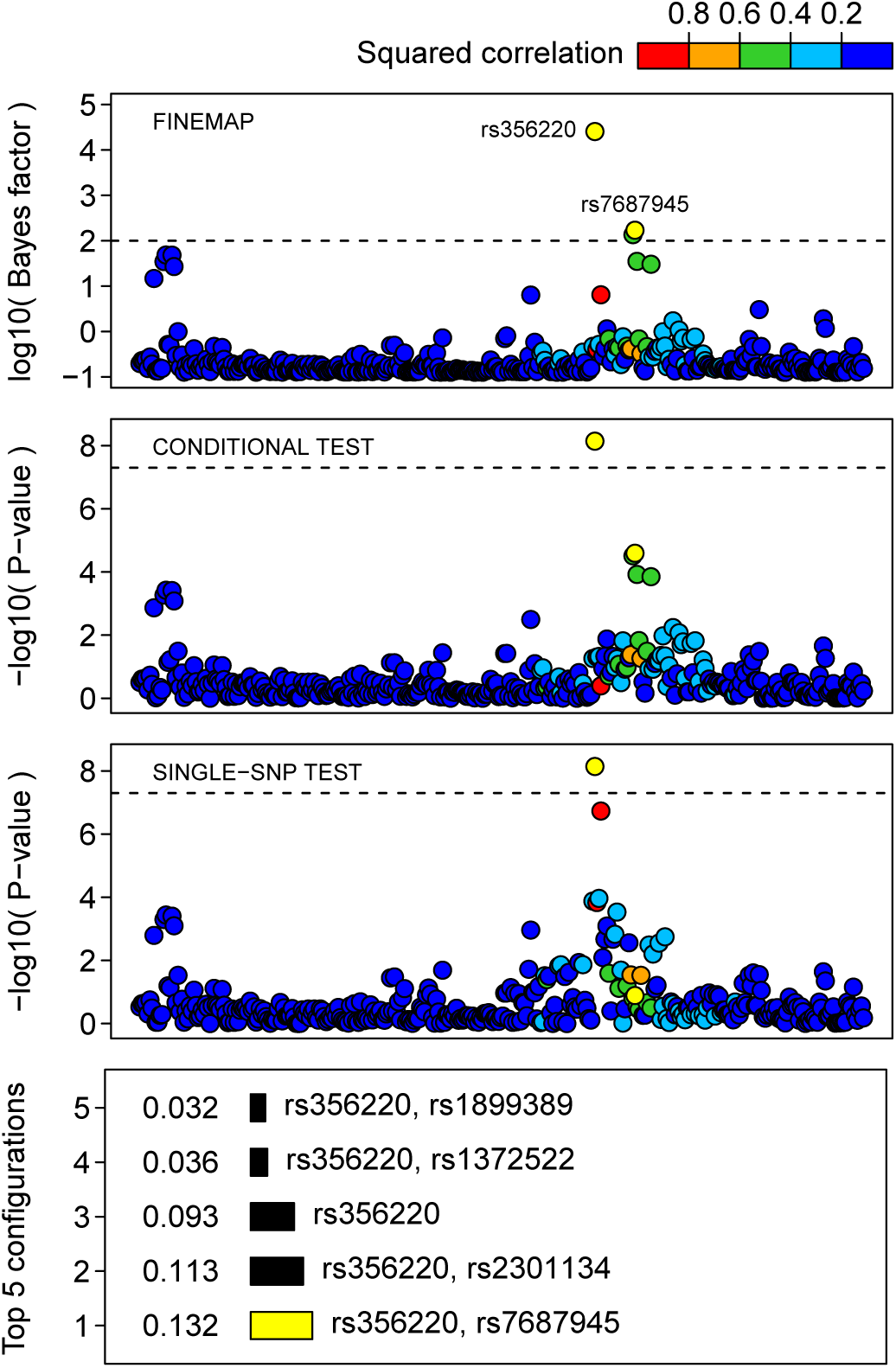
Fine-mapping of 4q22/SNCA region associated with Parkinson's disease. Associated SNPs rs356220 and rs7687945 are highlighted by 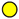 and their configuration by 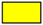. Dashed lines correspond respectively to a single-SNP Bayes factor of 100 and P-value of 5 × 10^−8^. Squared correlations are shown with respect to rs356220.

## 6 Discussion

GWAS have linked thousands of genomic regions to complex diseases and traits in humans and in model organisms. Fine-mapping causal variants in these regions is a high-dimensional variable selection problem complicated by strong correlations between the variables. We introduced a software package FINEMAP that implements an important solution to the problem: a stochastic search algorithm to circumvent computationally expensive exhaustive search. In all data sets we have tested, FINEMAP achieves similar accuracy as the exhaustive search but uses only a fraction of processing time. For example, fine-mapping a genomic region with 1500 SNPs allowing for at most 3 causal variants completes in 1.4 seconds using FINEMAP while the exhaustive search implemented in CAVIARBF requires about 5.2 hours. Computationally efficient algorithms are a key to handle ever increasing amount of genetic variation captured by emerging sequencing studies as well as to scale up the analyses to whole chromosomes or even to whole genomes.

FINEMAP uses a Shotgun Stochastic Search (SSS) algorithm (Hans *et al.*, 2007). SSS has been inspired by Markov Chain Monte Carlo (MCMC) algorithms that are widely used for Bayesian inference. For a review on MCMC, see Andrieu *et al.* (2003). Standard MCMC methods, such as the Metropolis-Hastings algorithm (Metropolis *et al.*, 1953; Hastings, 1970) and Gibbs sampler (Geman *et al.*, 1984), perform a sequence of steps in the parameter space via a stochastic transition mechanism that ensures a valid approximation to the target distribution. MCMC can often quickly reach an interesting region of the parameter space, but, at each step, it only considers one of the possible neighboring states. This means that MCMC is often slow to explore a high-dimensional state space. To improve on this, SSS generates a whole set of neighboring configurations at each iteration and saves them all for further use in probability calculations. This way a large number of parameter configurations with relatively high probability is quickly explored.

FINEMAP is accurate when the set of causal configurations explored captures a large majority of the total posterior probability. Our results show that this is the case in all data sets we have tested: the maximal error in any single-SNP inclusion probability is smaller than 0.11 across all 2000 data sets of Scenario B. Using exhaustive search, we observed in genomic regions with 750 SNPs of which five were truly causal that on average only the top 123 (median = 14) causal configurations out of all possible 70.3 × 10^6^ already cover 95% of the total posterior probability. (Similar results were also observed for genomic regions with different numbers of SNPs.) This explains why an efficient stochastic search can achieve accurate results in a tiny fraction of the processing time of an exhaustive search. Our data sets were generated by requiring that the causal SNPs had highly correlated proxies (absolute correlation greater than 0.5) among the other variants. The high accuracy of FINEMAP throughout these tests makes us believe that FINEMAP is accurate in typical GWAS data with complex correlation structure among the SNPs.

Although we have not encountered any data set where FINEMAP would not have performed well, theoretically, it remains possible that an in-exhaustive search could miss some relevant causal configurations. A simple way to assess possible problems is to run many searches in parallel and compare and combine their outcomes. Another way is parallel tempering (Geyer, 1991) where several searches are run in parallel in different ’’temperatures”. Intuitively, increasing temperature flattens the likelihood function and hence a search in a higher temperature moves around more freely than one in a colder temperature. Such an approach, together with complex global transition mechanisms to escape from local modes, was introduced in an evolutionary stochastic search algorithm by Bottolo *et al.* (2010) that was later tailored for genetic analyses of multiple SNPs and multivariate phenotypes in the software package GUESS (Bottolo *et al.*, 2013). These two papers could give ideas how FINEMAP could be further modified if trapping into local modes of the search space were encountered in real data analyses of GWAS regions.

The accuracy of FINEMAP depends on the quality of the SNP correlation estimate. For some populations, sequencing of many thousands of individuals have either already been carried out or will complete soon. This allows reliable fine-mapping in individual populations down to low-frequency variants with minor allele frequencies above 0.5%. A more challenging problem is large meta-analyses that combine individuals from varying ancestries. Assuming that the causal variants are included in the data and have the same effect sizes across the ancestral backgrounds, FINEMAP can be run with the sample size weighted SNP correlation matrix. If these assumptions are not met, then a hierarchical model allowing separate SNP correlation structures in each ancestry would perform better (Kichaev *et al.*, 2015).

The output from FINEMAP is a list of possible causal configurations together with their posterior probabilities and Bayes factors similar to CAVIARBF. These probabilities contain all the information from the model needed for downstream analyses. Examples of useful derived quantities are the single-SNP inclusion probabilities, single-SNP Bayes factors, credible sets of causal variants (WTCCC *et al.*, 2012) and a regional Bayes factor against the null model (Chen *et al.*, 2015). We believe that FINEMAP, or related future applications of shotgun stochastic search to GWAS summary data, provide new opportunities to reveal valuable information that could otherwise remain undetected due to computational limitations of the existing fine-mapping methods.

## Acknowledgement

This work was financially supported by the Doctoral Programme in Population Health (C.B.), the Academy of Finland [257654 and 288509 to M.P.; 251217 and 255847 to S.R.] and a Wellcome Trust Career Development Fellowship to C.C.A.S [097364/Z/11/Z]. S.R. was further supported by the Academy of Finland Center of Excellence for Complex Disease Genetics, EU FP7 projects ENGAGE (201413), BioSHaRE (261433), the Finnish Foundation for Cardiovascular Research, Biocentrum Helsinki, and the Sigrid Jusélius Foundation.

We thank the participants of the FINRISK cohort and its funders: the National Institute for Health and Welfare, the Academy of Finland [139635 to Veikko Salomaa] and the Finnish Foundation for Cardiovascular Research.

This study makes use of GWA data generated by the Wellcome Trust Case-Control Consortium 2 (WTCCC2) on UK PD cases and on UK controls from the 1958 Birth Cohort (58BC) and National Blood Service (NBS).

The authors also wish to acknowledge CSC - IT Centre for Science, Finland for computational resources.

## References

Andrieu, C et al. (2003) An introduction to MCMC for machine learning, Mach Learn, 50, 5–43.

Bottolo, L. and Richardson, S. (2010) Evolutionary stochastic search for Bayesian model exploration, Bayesian Anal, 3, 583–618.

Bottolo, L. et al. (2013) GUESS-ing polygenic associations with multiple phenotypes using a GPU-based evolutionary stochastic search algorithm, PLoS Genet, 9, e1003657.

Carbonetto, P. and Stephens, M. (2012) Scalable variational inference for Bayesian variable selection in regression, and its accuracy in genetic association studies, Bayesian Anal, 1, 73–108.

Chen, W. et al. (2015) Fine mapping causal variants with an approximate Bayesian method using marginal test statistics, Genetics, 200, 719–736.

Geman, S and Geman, D. (1984) Stochastic relaxation, Gibbs distributions, and the Bayesian restoration of images, IEEE Trans Pattern Anal Mach Intell, 6, 721–741.

Geyer, C. J. (1991) Computing science and statistics: proceedings of the 23rd symposium on the interface, 156–163.

Guan, Y. and Stephens, M. (2011) Bayesian variable selection regression for genome-wide association studies and other large-scale problems, Ann Appl Stat, 5, 1780–1815.

Hans, D. et al. (2007) Shotgun stochastic search for “large p” regression, J Am Stat Assoc, 102, 507–516.

Hastings, W. (1970) Monte Carlo sampling methods using Markov chains and their applications, Biometrika, 57, 97–109.

Hormozdiari, F. et al. (2014) Identifying causal variants at loci with multiple signals of association, Genetics, 198, 497–508.

Kichaev, G. et al. (2014) Integrating functional data to prioritize causal variants in statistical fine mapping studies, PLoS Genet, 10, e1004722.

Kichaev, G. and Pasaniuc, B. (2015) Leveraging functional-annotation data in trans-ethnic fine-mapping studies, Am. J. Hum. Genet., 97, 260–271.

Metropolis, N et al. (1953) Equations of state calculations by fast computing machines, J Chem Phys, 21, 1087–1092.

Pirinen, M et al. (2013) Efficient computation with a linear mixed model on large-scale data sets with applications to genetic studies, Ann Appl Stat, 7, 369–390.

Servin, B. and Stephens, M. (2007) Imputation-based analysis of association studies: candidate regions and quantitative traits, PLoS Genet, 3, e114.

Spain, S. and Barrett, J. (2015) Strategies for fine mapping complex traits, Hum. Mol. Genet., 42, 1001–1006.

The UK Parkinson’s Disease Consortium and The Wellcome Trust Case Control Consortium 2. (2011) Dissection of the genetics of Parkinson’s disease identifies an additional association 5’ of SNCA and multiple associated haplotypes at 17q21, Hum. Mol. Genet., 20, 345–353.

Vartiainen, E. et al. (2010) Thirty-five-year trends in cardiovascular risk factors in Finland, Int J Epidemiol, 39, 504–518.

Wallace, C. et al. (2015) Dissection of a complex disease susceptibility region using a Bayesian stochastic search approach to fine mapping, PLoS Genet, 11, e1005272.

Wellcome Trust Case Control Consortium et al. (2012) Bayesian refinement of association signals for 14 loci in 3 common diseases, Nat. Genet., 44, 1294–1301.

Wood, A. et al. (2014) Defining the role of common variation in the genomic and biological architecture of adult human height, Nat. Genet., 46, 1173–1186.

Yang, J et al. (2011) GCTA: a tool for genome-wide complex trait analysis, Am. J. Hum. Genet., 88, 76–82.

